# An improved demand curve for analysis of food or drug consumption in animal experiments

**DOI:** 10.1101/765461

**Authors:** Mark Newman, Carrie R. Ferrario

**Affiliations:** Center for the Study of Complex Systems, University of Michigan, Ann Arbor, Michigan, USA; Department of Pharmacology, University of Michigan, Ann Arbor, Michigan, USA

## Abstract

The incorporation of microeconomics concepts into studies using preclinincal self-administration procedures has provided critical insights into the factors that influence consumption of a wide range of food and drug reinforcers. In particular, the fitting of demand curves to consumption data provides a powerful analytic tool for computing objective metrics of behavior that can be compared across a wide range of reward types and experimental settings. The results of these analyses depend crucially on the mathematical form used to fit the data. The most common choice is an exponential form proposed by Hursh and Silberberg, which is widely used and has provided fundamental insights into relationships between cost and consumption, but it also has some disadvantages. In this paper we first briefly review the use of demand curves to quantify the motivating effects of food and drugs, then we describe the current methodology and highlight some potential issues that arise in its application. To address these issues, we propose a new mathematical framework for the analysis of consumption data, including a new functional form for the demand curve. We show that this proposed form gives good fits to data on a range of reinforcers across different animals and different experimental protocols, while allowing for straightforward calculation of key metrics of demand, including preferred consumption level, maximum response, price at maximum response, and price elasticity of demand. We provide software implementing our entire analysis pipeline, including data fits, data visualization, and the calculation of demand metrics.

## 1 Introduction

Behavioral economics is a branch of microeconomic theory and practice that aims to understand and quantify the economic choices made by individuals (Cartwright 2018; Kahneman 2011; Ayers and Collinge 2004; Perloff 2016). Classically, the field has focused on choices made by humans, but behavioral economic concepts have also been fruitfully applied to animal behavior, particularly in the context of self-administration procedures (Hursh 1993; Winger et al. 2002; Hursh et al. 2005; Winger et al. 2006; Hursh and Silberberg 2008; Galuska et al. 2011; Bentzley et al. 2013; Kawa et al. 2016; Pantazis et al. 2019). Although animals do not have an economy in the sense of markets, currency, credit and so forth, in many cases they are willing to “pay” for goods such as food or drugs by performing work. By measuring the amount of work they are willing to do for a given return one can develop notions of “price” and “demand” for goods, on which the tools of economics can then be brought to bear.

The utility of economic concepts in the study of food and drug motivation and addiction has been demonstrated extensively since their introduction to the field in the 1990s (Hursh 1993; Hursh and Winger 1995; Bickel et al. 2000; Hursh and Silberberg 2008). Foundational contributions by Hursh, Winger, Bickel, and others have established behavioral economics approaches as a fundamental tool for the quantitative assessment of self-administration behavior. The approach has been widely applied to evaluate motivation across different reinforcers or testing conditions, to study effects of pharmacological or behavioral interventions on demand, and to address abuse liability and neurobehavioral underpinnings of substance abuse.

The behavioral economics approach as currently applied focuses on the study of the so-called “demand curve,” which represents the consumption of food or drug as a function of the work or “price” required to obtain it. Such curves are computed by fitting an appropriate mathematical form to raw self-administration data, and aspects of the fitted curve provide summary statistics describing features such as the maximum effort exerted to obtain a given reinforcer and the price at which that maximum effort occurs. Most previous work has made use of an exponential form for the demand curve proposed by Hursh and Silberberg (2008) but, as we discuss, this form and the accompanying methodology have some shortcomings that create challenges for comparisons across studies and for reproducibility of results.

In this paper, we present a new approach for fitting self-administration demand data that addresses these shortcomings while providing easy calculation of key metrics of preferred consumption, price sensitivity, and motivation. We begin with a brief introduction to the relevant concepts of behavioral economics followed by a description of the current methodology. We then describe our proposed approach, outlining its main features and providing a simple recipe for data analysis that gives straightforward quantitative answers to questions relevant to the study of demand, motivation, and addiction.

## 2 Economics of consumption

Consider a typical self-adminstration experiment in which an animal is given the opportunity to perform work, such as lever presses, in return for a desirable good. In most cases the good is either food or drug; for the purposes of illustration let us say it is a dose of drug. The “price” of the drug can be varied by the experimenter, usually using one of two methods: either they can vary the number of lever presses required to receive a fixed dose, or they can vary the dose received for a fixed number of lever presses. Either way, one can define the price *P* of the drug as the number of lever presses per unit of drug received, measured for instance in milligrams. Thus:

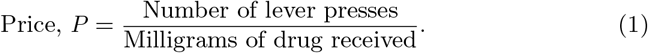

(See Table 1 for a summary of the variables used in this paper.)

**Table 1.** Summary of the variables used in the theory described here, along with alternative notations for the same quantities used by other authors.

| Variable | Meaning | Other notations |
| --- | --- | --- |
| $P$ | Price, work performed per unit of good received | $C$ |
| $Q$ | Consumption | |
| $R$ | Revenue, total work performed at a given price point | $O$ |
| $Q_0$ | Preferred consumption level when price is negligible | |
| $R_{\max}$ | Maximum work performed to obtain goods at any price | $O_{\max}$ |
| $P_{\max}$ | Price at which maximum work is performed | |
| $\tilde{P}_{\max}$ | Normalized $P_{\max}$, equal to $Q_0 P_{\max}$ | $P_{\max}, nP_{\max}$ |
| $\alpha$ | Fitted parameter in exponential model, inversely proportional to $P_{\max}$ | |
| $\tilde{\alpha}$ | Normalized $\alpha$, equal to $\alpha/Q_0$, also called “essential value,” inversely proportional to $\tilde{P}_{\max}$ | $\alpha$ |

In the most straightforward version of the experiment, the experimenter allows the animal to “buy” repeated doses of drug at a set price and records the total amount of drug *Q* consumed during a session of fixed length (anywhere from a few minutes to hours, depending on the drug, the question at hand, and so forth). The procedure can then be repeated for a range of different prices to measure consumption *Q* as a function of price *P*.

The data produced by experiments of this kind have a characteristic form. First, there is normally a clear preferred amount of drug that a particular animal will consume in the allowed time when price is not an issue. Even if the price is reduced practically to zero so that drug is essentially free, the animal will not consume an unlimited amount, but will stop when it reaches its preferred level of consumption. Traditionally this level is denoted *Q*_0_.

Next, if we now raise the price slightly, so that drug is not free but still very cheap, the animal will still be willing to do the modest work required of it and will take its fill of drug, meaning its consumption will still be *Q*_0_. But if we raise the price enough, the effort will start to become a factor and the animal will consume less drug. And if the price is very large—if the animal has to do a million lever presses, say, to receive an infusion—then consumption must be zero, since it is physically impossible to perform this many lever presses in the allowed amount of time.

Thus, when plotted against price, we expect consumption to look something like Fig. 1a. The data points in this figure show actual consumption against price for a rat self-administering cocaine.^1^ Observe how the points are roughly flat in the left part of the plot, but fall off beyond a certain “breakpoint,” denoted roughly by the vertical dashed line, as the price rises. If one were to continue the measurements far enough to the right of the plot, they would eventually reach zero when the price becomes so high that the rat receives no drug at all. Note that the graph is plotted on logarithmic scales, a standard practice that allows us to capture the typically wide range of values of both price and consumption.

**Fig. 1.**
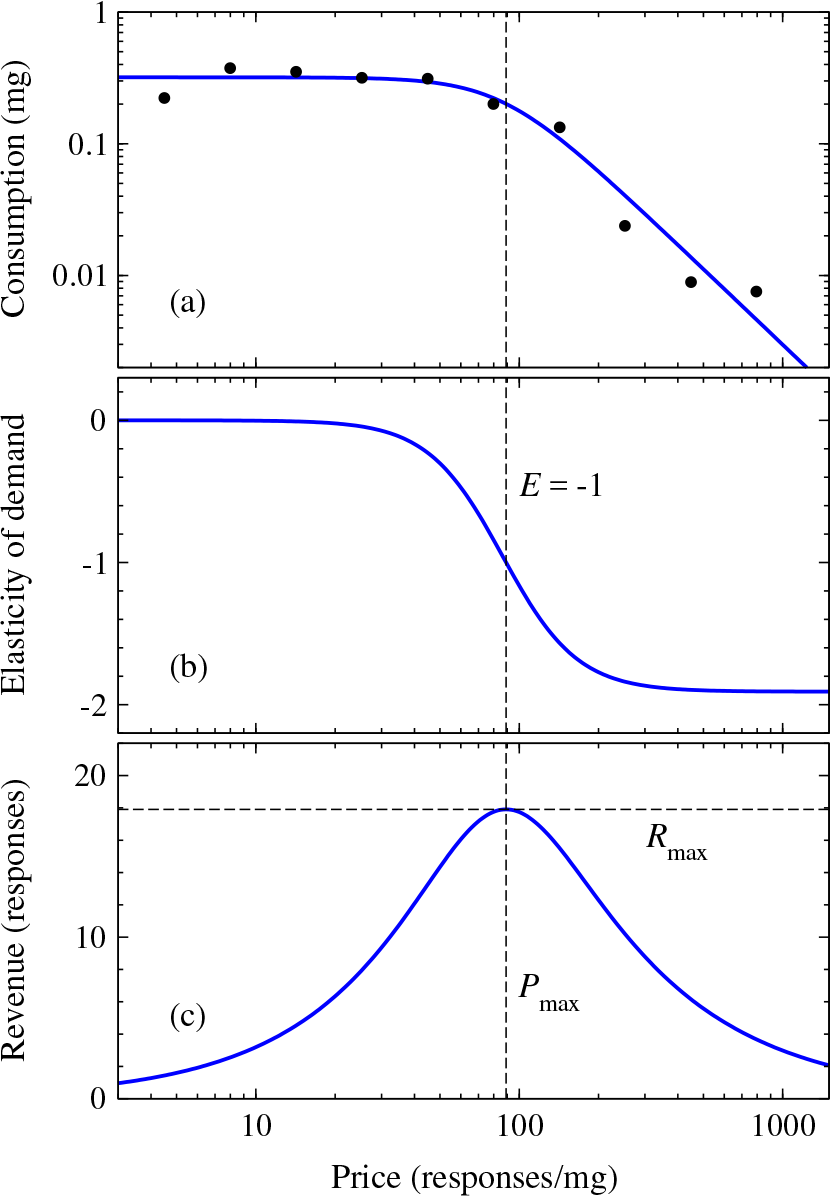
(a) Data points: total consumption of self-administered cocaine by a single male rat as a function of price *P* measured in lever presses per milligram. Solid curve: the demand curve reconstructed by fitting to Eq. (21). (b) The elasticity corresponding to the demand curve in (a). Note that the value of the elasticity is always negative and that the point where elasticity equals 1 coincides with the point at which the revenue [plotted in (c)] reaches its maximum. (c) The “revenue” corresponding to the demand curve in (a), i.e., the total work performed at each price point. The maximum revenue *R*_max_ falls at price *P*_max_, which coincides with the point at which elasticity is −1.

## 3 The demand curve

Data of the kind shown in Fig. 1a can be summarized by fitting a *demand curve* to it. An example is shown as the solid curve in Fig.1a; it represents the expected consumption level *Q* as a function of price *P*. The demand curve has become a standard tool in the analysis of consumption data, capturing in a single graph the willingness of an animal to work for a range of outcomes. The utility of this approach has been demonstrated repeatedly and demand curve analyses have provided essential information about many aspects of consumption and drug-taking behavior in both humans and animal models [see for example Hursh and Silberberg (2008); Bentzley et al. (2013); Aston and Cassidy (2019)]. As we will see, the demand curve allows us to describe motivation for drugs or food in a quantitative manner, placing numbers on concepts that may otherwise be accessible only via more qualitative approaches, and allowing comparisons across different reinforcers [see Hursh and Winger (1995) and Hursh et al. (2005) for reviews]. To do this we borrow some concepts from economics, starting with the so-called elasticity of demand.

### 3.1 Elasticity of demand

Suppose that we know the demand curve for a particular experiment—the solid line running through the data points in Fig. 1a. (We will see shortly how to extract such curves from data.) In general the demand curve falls off as price increases, since we expect an animal to consume less of a good as the work required to obtain it increases (the so-called “law of demand”). The *price elasticity of demand E* measures exactly how the demand curve falls off with increasing price. For instance, if we double the amount of work a rat has to do to receive a dose of drug, will the rat consume the same amount of drug overall? Half as much? A quarter?

The elasticity is the ratio between the fraction the price goes up by and the fraction the consumption goes down by. For example, if price goes up by 10% and as a result consumption falls by −20%, then the elasticity is *E* = −20*/*10 = −2. Note that elasticity is normally a negative number, as here.

More generally, suppose that the price *P* increases by an amount *dP*. Then the *fraction* that price increases by is *dP/P*. If at the same time consumption *Q* goes down by *dQ* then the fractional decrease in consumption is *dQ/Q*. The elasticity is the ratio of these two fractions (fractional decrease in consumption over fractional increase in price), which is

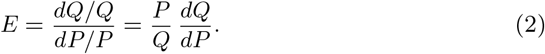

The quantity *dQ/dP* is the derivative of consumption with respect to price.^2^ The derivative is the slope of a graph of *Q* against *P*. Because the graph is downward sloping in this case, the slope is negative, and hence again *E* will be a negative number.

Alternatively, we can note that

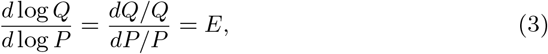

which is the same elasticity again, meaning that *E* is also equal to the slope of a graph of log *Q* against log *P*. In other words, if we plot the demand curve on log scales (as in Fig. 1a), then the elasticity is the slope.

Figure 1b shows the elasticity for the demand curve of Fig. 1a. Note that in general elasticity is not a single number: it varies with price. In the left part of Fig. 1a, for instance, the demand curve is flat and hence the elasticity in Fig. 1b is close to zero. But in the right part the demand curve slopes downward quite steeply, meaning the elasticity is large and negative.

The elasticity is widely used in (human) economics as a measure of the price sensitivity of goods, including psychoactive substances (Ayers and Collinge 2004; Perloff 2016). A large (negative) elasticity of demand indicates a good that is highly price sensitive: small increases in price will substantially decrease demand. A small elasticity indicates a relatively price-insensitive good. For instance, the elasticity of soft drinks has been measured to be about −3.8 at prevailing prices (Ayers and Collinge 2004), indicating substantial price sensitivity—if the price of a soft drink is increased people will simply stop drinking it. On the other hand, the elasticity of cigarettes is estimated to be much smaller, around −0.4 (Becker et al. 1994), indicating significantly lower price sensitivity—people will continue to smoke even if the price of cigarettes goes up.

These results suggest that elasticity could be used as a measure of the reinforcing or motivating effects of drugs or food, not only in humans but also in animal experiments. In practice, however, it is rarely used in this way in the animal literature. Instead, elasticity has primarily been of interest because of its role in estimating the maximum work that animals perform, as we discuss in the next section. (There are claims in the literature of using elasticity to quantify motivation, but in most such cases no value of the elasticity is actually reported. There appears to be some misunderstanding of what elasticity is and what it represents, as we discuss in Section 4.2.)

### 3.2 Measuring motivation

One of the primary uses of demand curves in animals is for quantifying responding for food or drug. How motivated are animals to take a drug? Can we define a single number to quantify motivation? How does motivation change over time or in relation to other reinforcers? One approach is to look at the total amount of work an animal is willing to perform over the course of an experimental session. In the jargon of economics this total amount of work is called the *revenue*, denoted *R*, although in the present context it may be more useful to think of the *R* as standing for “responses,” since it is simply equal to the number of lever presses or other work the animal performs.^3^

We have seen that the price *P* is defined as the amount of work that must be performed to receive one unit of the desired good, such as the number of lever presses per milligram of drug [see Eq. (1)]. If we know the number of lever presses per milligram and we also know the total number of milligrams consumed *Q*, then the total number of lever presses—the revenue *R*—must be the product of the two:

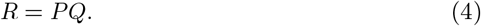

Figure 1c shows a plot of the revenue corresponding to the demand curve in Fig. 1a. Observe how the revenue starts at a low value on the left of the plot in the regime where the animal is required to perform only a little work to receive drug, rises to a maximum around the “breakpoint” in the demand curve, then falls off again as the price becomes too high and the animal abandons trying to obtain drug. These observations suggest two possible measures of the motivation potential of food or drugs: (1) the maximum amount *R*_max_ of work—the maximum revenue—the animal is willing to commit to obtaining goods at any price,^4^ or (2) the price *P*_max_ at which this maximum occurs. In Fig. 1c, *R*_max_ corresponds to the height of the peak in revenue (the horizontal dashed line) and *P*_max_ corresponds to the price at which that peak occurs (the vertical dashed line). Perhaps more intuitively, *P*_max_ corresponds roughly to the breakpoint in the demand curve, Fig. 1a, at which the curve falls off from its initial plateau. Thus *P*_max_ measures the maximum price the animal will tolerate before it gives up and reduces its consumption.

To calculate *P*_max_ we maximize Eq. (4) with respect to *P* by differentiating thus:

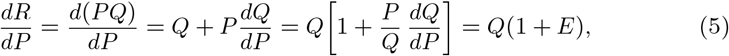

where *E* is the elasticity as before and we have used Eq. (2). Setting Eq. (5) to zero we then find that the maximum revenue is achieved when the elasticity *E* is equal to −1, and thus we can find *P*_max_ by finding the price at which this occurs. We give examples of this calculation in Section 4.4 and 5.2. Once we have determined *P*_max_ we can substitute the result back into Eq. (4) to find the corresponding value of the revenue, *R*_max_.

Ease of calculating *P*_max_ and *R*_max_ is one of the reasons why the demand curve and the elasticity are useful concepts. In principle, one could imagine measuring *P*_max_ and *R*_max_ directly from the data by asking what the maximum number of responses is at any price and at what price that maximum falls. Ad hoc methods for doing this have been proposed, for instance, by Hursh and Silberberg (2008) [see also Oleson et al. (2011)]. The results returned by these methods, however, are limited to the specific values of price and responding measured in the experiment and so give only a general indication, and they are moreover prone to measurement fluctuations and hence can be unreliable. Calculations based on demand curves are more robust and repeatable (Bentzley et al. 2013).

Both *R*_max_ and *P*_max_ are reasonable measures of motivation, but they are not equivalent. If two animals display the same *P*_max_ for a given reinforcer but the first has higher *R*_max_ it implies that the first is willing to do more work than the second for the same amount of consumption. Conversely, if they display the same *R*_max_ but the first has higher *P*_max_ then the first is willing to do the same amount of work for less consumption. Certainly these two measures could be correlated, but they are not the same thing.

*R*_max_ has found use in human studies, where strong associations have been observed between its value and, for instance, post-intervention alcohol consumption (MacKillop and Murphy 2007; MacKillop et al. 2009), although it should be noted that there were also strong relationships in these studies among *P*_max_, *R*_max_, and other demand metrics (see Section 6 and Fig. 6 for further discussion). Hursh and Winger (1995) make a persuasive argument that *R*_max_ is essentially independent of the potency or magnitude of a reinforcer, which argues in favor of its use for comparisons between different reinforcers. The value of *P*_max_, by contrast, varies with potency and hence *P*_max_ appears inferior in this respect. However, it is possible by suitable normalization to create a potency-independent version of *P*_max_, as discussed in Section 3.3. Moreover, *R*_max_ is difficult to measure in some situations, particularly when using progressive ratio schedules, whereas *P*_max_ is relatively easy to measure (Richardson and Roberts 1996). Thus both *P*_max_ and *R*_max_ are useful in certain contexts, and both are employed in the field, although arguably *P*_max_ is more common.

### 3.3 Normalized price and comparisons between different reinforcers

In addition to their use in basic data analysis, demand curves are used as a way to compare behavior across experiments on different drugs or other reinforcers. Can we tell, for instance, whether animals have greater motivation for food or drugs? Or for one drug over another?

One way to perform such comparisons is to use the measure *R*_max_ defined in Section 3.2 above, which is the maximum work animals are willing to perform to obtain a good, at any price. Hursh and Winger (1995) argue that *R*_max_ is well suited to comparisons between different reinforcers and describe it as “a sensitive tool for direct comparison and quantitative ordering of demand, both within and across the drug classes (stimulant, sedative, and opioid)”.

Conversely, the measure *P*_max_ (also defined in Section 3.2), which is the price at which animals exert their greatest effort to obtain goods, is not well suited to answering such questions because it is not clear how one should compare prices for different goods. Is a price of 10 lever presses per milligram of cocaine higher or lower than 10 lever presses per milligram of amphetamine? The answer depends on the potency of the drugs in question: one milligram might have a strong effect for one drug but only a weak effect for another. To make a meaningful comparison, we need to normalize the price by a suitable factor that represents the typical magnitude of drug intake in milligrams (or other suitable units). Fortunately, we have exactly such a factor to hand, namely the preferred consumption level *Q*_0_.

If we divide dose by *Q*_0_ we get a number that is independent of potency: every dose is specified as a fraction of the preferred consumption level for the same good. Thus, a suitable normalized price, as first proposed by Hursh and Winger (1995), is given by replacing the dose in Eq. (1) by dose divided by *Q*_0_:

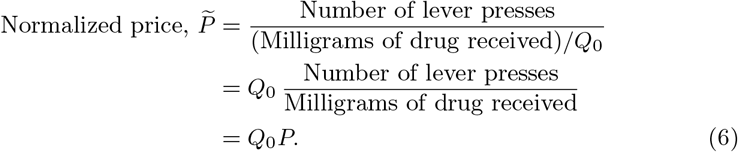

Combining this approach with our measure *P*_max_, we can then write a potency-independent measure of motivation thus:

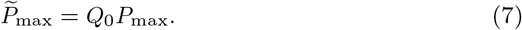

This measure appears to work well in practice for comparing demand across different reinforcers.

Equations (6) and (7) are not in precisely the form used elsewhere. In much of the literature, for instance, the price is denoted *C* instead of *P* and the normalized price *P* is denoted *P*. This, however, is merely a matter of notation. A more substantive difference is that many experimenters avoid the use of *P*_max_ as a measure of motivation in favor of another parameter commonly denoted *α*. We discuss *α* in detail below (Section 4.4), where we show that it is in fact essentially equivalent to *P*_max_ but has some disadvantages that *P*_max_ does not share.

## 4 Fitting the demand curve

Data such as those shown in Fig. 1a already give us a rough outline of the demand curve. But they also inevitably display statistical fluctuations and moreover give us the consumption at only a small set of discrete price points. We can reduce the effects of fluctuations and interpolate between price points by fitting a suitable curve through the data, recovering the entire demand curve, as shown by the blue line in Fig. 1a. The fitting procedure itself is straightforward—there are a range of software packages that will do the job. A crucial question, however, is what mathematical form the fitted curve should take. We need to specify a form that is flexible enough to fit the data we see in experiments on a range of different consumables, different animals, and different procedures, while at the same time following the common-sense requirements that the curve be flat at first, then drop off, and go to zero as price becomes large.

### 4.1 The exponential demand curve

The most common mathematical form used for demand curves in self-administration experiments of the kind considered here is the “exponential” form advanced by Hursh and Silberberg (2008):

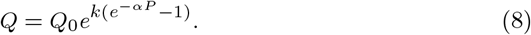

This equation relates the consumption *Q* to the price *P* using three parameters. The first is *Q*_0_, which we have already discussed—it is the preferred level of consumption when the price is low enough to have no limiting effect on intake. The other two parameters are *k* and *α*, which we look at more closely in the following section.

Because demand curves are normally plotted on logarithmic scales, one often sees Eq. (8) expressed in terms of the logarithm of *Q*. Taking the log of both sides of the equation we find that

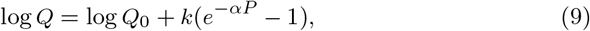

where “log” denotes the natural logarithm (base *e*).^5^ The two forms, Eqs. (8) and (9), are entirely equivalent and contain the same information.

### 4.2 Parameters for the exponential demand curve

Though it has been widely employed, the demand curve defined in Eq. (8) has some shortcomings. Specific issues include difficulty estimating or interpreting the parameters *k* and *α*, difficulty estimating the elasticity, and an unrealistic nonzero value of consumption at large prices.

#### The parameter k and the limiting value of Q

A disadvantage of Eq. (8) is that it fails to meet one of our fundamental criteria for a demand curve, that it go to zero as price becomes large. As we have said, it is axiomatic that the curve should go to zero: if the price of drug is a million lever presses per dose then the animal is necessarily going to consume no drug. As *P* goes to infinity, however, the form in Eq. (8) tends to the limiting value *Q*_0_*e*^−*k*^, which is always nonzero.

This conflict causes a number of problems. First, it is undesirable to a fit curve to data when we know the curve to have a different shape from the data. Just as one should not fit a straight line if one knows the data to follow a rounded form, so one should not fit data that must go to zero with a form that does not. This is, however, a somewhat theoretical objection. A more practical issue arises when we attempt to estimate the parameter *k*. Since *k* controls the value of consumption when price becomes large, one should be able to determine *k* by measuring this value. This, however, is not possible in the present case, since as we have said there is no such value in practice: consumption always goes to zero.

There are ways around this difficulty. One could for instance estimate *k* using some other type of fitting procedure. Hursh and Silberberg (2008) take a different route, avoiding the problem altogether by not fitting *k* to the data at all. Instead, they choose its value themselves, writing that “The value of *k* is generally set to a common constant across comparisons because it merely specifies the range of the data.” In related work Bentzley et al. (2013) write that the value of *k* is “chosen based on the maximum observed range of consumption.” Specifically, they compute the range spanned by the observed values of log *Q* for all sessions and set *k* equal to the largest such range. Gilroy et al. (2019) suggest a slight variant of this procedure, calculating the same maximum range but then adding 0.5, to guard against the possibility that *k* “does not reflect the full range of observed consumption values.”^6^

In practice, however, approaches such as these are somewhat unsatisfactory because they determine *k* using an ad hoc recipe rather than by fitting to the data. Such recipes can result in different experimenters using different values of *k*— and hence reaching different conclusions—even when fitting to the same set of data points. Consider Fig. 2, which shows the same cocaine self-administration data that appears in Fig. 1a. The four curves in the figure show the best fit of the exponential demand curve to these data for four different values of *k*. The blue (solid) curve shows the fit when *k* is chosen according to the prescription of Bentzley et al. that *k* should be set to the largest range spanned by log *Q* over all sessions. Since we are looking at only a single session in this case, *k* is simply equal to the range of the data, which gives a value of about 4. (The exact value is 3.91, but we use 4 for simplicity.)

**Fig. 2.**
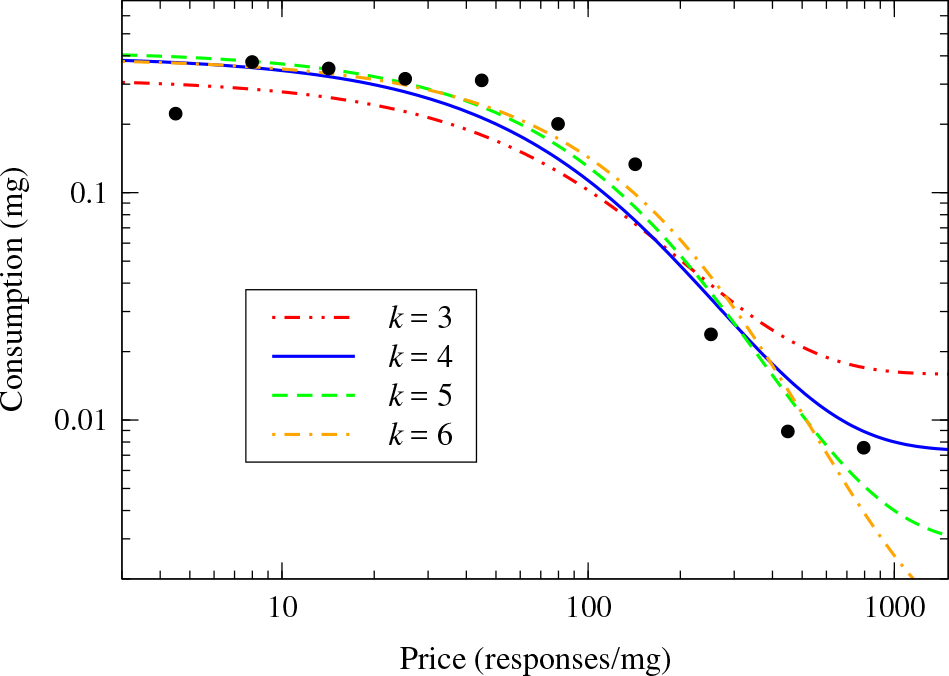
Best fits of the cocaine self-administration data from Fig. 1a to the exponential demand curve of Eq. (9) for various choices of the parameter *k* as indicated.

On the other hand, if we were examining these data as one session out of many, it is likely that at least one other session would have a larger range of log *Q*, meaning that we would have to use a larger value of *k*. The green and orange (dashed and dot-dashed) curves in Fig. 2 show fits with *k* = 5 and 6. For comparison we also show one fit with a smaller value of *k* = 3 (red, dot-dot-dashed).

The values of the parameters of the fit for each curve are shown in Table 2. As we can see, the values span quite a wide range. The value of *Q*_0_ varies from 0.319 to 0.420 for example, an increase of 32%, and *P*_max_ shows a similar increase of 38%. Also shown in the table is the quantity we call 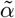, which is described in Section 4.4 and which is widely used as a measure of motivation in the literature. The fitted values of this quantity vary over a broad range from a low of 0.0047 to a high of 0.0149, an increase of 217%. The quantity 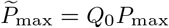, which we recommended in Section 3.3 as a measure of motivation, fares better, varying from 37.5 to 43.6, an increase of just 16%, but even this variation is large enough to inject significant uncertainty into the results, given that it is provoked solely by making different choices for the parameter *k*.

**Table 2.** Values of the parameters *Q*_0_ and *α* for the four fitted curves in Fig. 2, each for a different value of *k* as indicated, along with the calculated values of the quantities 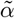, *P*_max_, 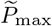, and *R*_max_.

| $k$ | $Q_0$ | $\alpha$ | $\tilde{\alpha}$ | $P_{\max}$ | $\tilde{P}_{\max}$ | $R_{\max}$ |
| --- | --- | --- | --- | --- | --- | --- |
| 3 | 0.319 | 0.00476 | 0.01491 | 130.2 | 41.5 | 10.4 |
| 4 | 0.400 | 0.00379 | 0.00948 | 94.3 | 37.7 | 11.3 |
| 5 | 0.420 | 0.00267 | 0.00636 | 96.9 | 40.7 | 13.0 |
| 6 | 0.390 | 0.00183 | 0.00469 | 111.8 | 43.6 | 14.4 |

#### The parameter α

The parameter *α* is also somewhat problematic, although for different reasons. In principle this parameter is unexceptionable: it plays the role of an exponential constant and can be estimated in straightforward fashion by fitting to data such as those in Fig. 1a. The problem is that in much of the literature *α* is said to be a measure of elasticity and it has been extensively treated as such [e.g., Koffarnus et al. (2011); Bentzley et al. (2013); Lacy et al. (2019)]. As discussed in Section 3.1, elasticity is a well-defined and useful concept in economics, but *α* is not a measure of elasticity. Rather, *α* is a measure of the price at which the animal performs maximum work, equivalent to the quantity *P*_max_ discussed in Section 3.2 above. This point is made clearly in the literature, but at the same time misinterpretation of *α* is also common. For example, Bentzley et al. (2013) state correctly that “*α* is an inherently normalized parameter and equivalent to [normalized *P*_max_], as these variables are inversely proportional”, but also that “*α* is a measure of demand curve elasticity.” To some extent this may be a matter of semantics—the scientific conclusions are largely unaffected—but in the interests of clarity it is good to be precise about the role of the variables.

A further issue with *α* is that its value is strongly influenced by the choice of *k*. In Sections 4.3 and 4.4 we examine the elasticity for the exponential demand curve in detail and explain the role played by *α*.

### 4.3 Elasticity for the exponential demand curve

Another issue with the exponential demand curve arises when we attempt to estimate the corresponding elasticity of demand. The elasticity for Eq. (8) can be calculated by taking the derivative of the logarithmic form in Eq. (9), which, as shown by Bentzley et al. (2013), gives

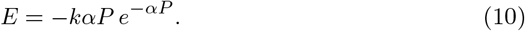

Unfortunately, this value depends fundamentally on the parameter *k* which, as we have said, is not determined from the data, but instead is chosen by the experimenter. To see an example of why this matters, let us calculate the maximum value of the elasticity *E*_max_ for given values of the parameters *Q*_0_, *α*, and *k*. To do this we differentiate (10) with respect to *P* and set the result to zero giving:

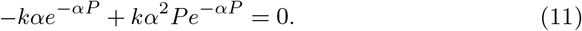

Canceling a number of factors and rearranging, we find that *αP* = 1, and substituting back into (10) we find the maximum value of the elasticity to be

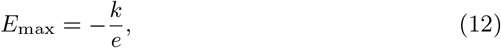

where *e* = 2.718 … is the base of the natural logarithm.

Thus the maximal value of the elasticity depends only on *k* (and the mathematical constant *e*). Since *k* is determined using an ad hoc recipe that can give different answers for the same data depending on experimental context, this means that the value of *E*_max_ is largely arbitrary.

The net result is that in most cases values of the elasticity determined from data fits to Eq. (8) [or Eq. (9)] are not informative. This may be one reason why elasticity has not found wide use in the analysis of self-administration data.

### 4.4 Maximum work performed for the exponential demand curve

We have seen that the price *P*_max_ at which maximum work is performed falls at the point where the elasticity *E* is equal to −1 [Section 3.2 and Eq. (5)]. For the exponential demand curve, the elasticity is given by Eq. (10) and hence maximum work occurs when

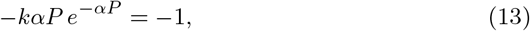

which can be solved to give

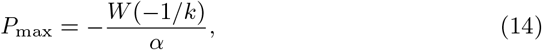

where *W* is the so-called Lambert *W*-function (Olver et al. 2010; Gilroy et al. 2019).^7^ Once we have the value of *P*_max_ it is straightforward to compute the corresponding value of *R*_max_ by substituting into Eqs. (4) and (8), which gives

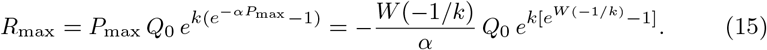

In current approaches to data analysis, the standard procedure is to hold the value of *k* constant over different sessions and different animals, meaning that *W* (−1/*k*) is also constant so that Eq. (14) implies *P*_max_ is inversely proportional to *α* and hence, as discussed in Section 4.2, the two quantities measure the same thing, a point that has been made clearly by, for example, Bentzley et al. (2013). The detailed relationship between *α* and *P*_max_ however still depends on the value of *k* since Eq. (14) can be rearranged to read

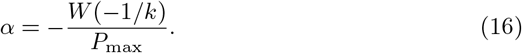

This means that even when working with identical data, from animals with the exact same *P*_max_, different experimenters will arrive at different results for *α* if they use different values for *k*. Under the circumstances, therefore, we do not recommend using the value of *α* as a measure of motivation, given that *P*_max_ itself contains the same information but is not directly dependent on *k*.

Some writers have argued in favor of using *α* because for some formulations of the demand curve it automatically incorporates a normalization by a factor of *Q*_0_ of the kind discussed in Section 3.3. One can rewrite the demand curve of Eq. (9) thus:

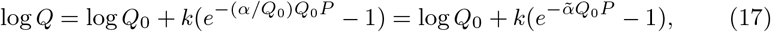

where

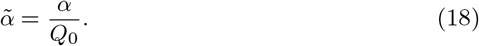

Equation (17) is exactly equivalent to the original form (9) but expresses the consumption as a function of the normalized price *Q*_0_*P* of Section 3.3. This is the form in which the exponential demand curve is mostly commonly written,^8^ and it involves a change in the definition of *α* according to (18). Combining (18) with Eq. (16), we find that

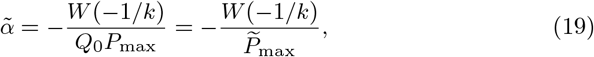

where we have used the definition of 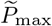 from Eq. (6).

In other words 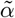 is inversely proportional to the normalized measure 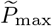 and hence measures the same thing. Because of this, 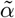 has been used widely as a measure of motivation and is in fact probably the most commonly used such measure. It is sometimes referred to as the “essential value,” and often denoted simply *α*, though we prefer the notation 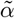 to avoid confusion with the original *α* parameter of Eq. (8). While 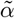 is an appropriate normalized measure of motivation, however, it still suffers from the same shortcoming as the unnormalized *α*, that it depends on the choice of *k*. We saw an example of this issue in Section 4.2 (see Table 2). A further reason to avoid the use of 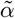 is that it is specific to the exponential demand curve form. The value of 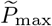 can be calculated for any demand curve, but 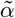, by its nature as a parameter of the exponential form, can only be calculated if one uses that form.

## 5 An alternative form for the demand curve

In Section 4 we examined the use of fitted demand curves as a way of quantifying the variation of consumption with price and discussed the widely used exponential form, Eq. (8), which has played an important role in the field but has some disadvantages—in particular that it does not go to zero as price becomes large, and that it depends crucially on the parameter *k* which is not determined by a fit to the data. Here we propose an alternative form for the demand curve which behaves in many ways like the exponential form but eliminates these shortcomings. The form we propose is also mathematically simpler, making solution of the resulting equations more straightforward. In the accompanying materials we provide an open-source data analysis program that calculates demand curves and parameters such as *P*_max_ and *R*_max_ from experimental data using our proposed form, as well as giving visualizations of the demand and revenue curves and best-fit parameter values.^9^ Instructions for using the program are given in the appendix.

### 5.1 Form of the proposed demand curve and parameters

The functional form we propose for the demand curve is

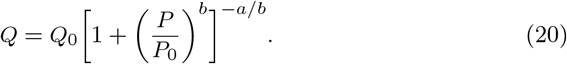

The parameters, depicted in Fig. 3, are as follows:

*Q*_0_: the preferred consumption level, as previously, i.e., the height of the plateau in the curve.
*P*_0_: the price at the breakpoint where the curve falls off, where the work is no longer worth the outcome.
*a*: the slope of the right-hand part of the curve where it falls off.
*b*: a parameter controlling the width of the “knee” or transition region between the left and right parts of the curve.

**Fig. 3.**
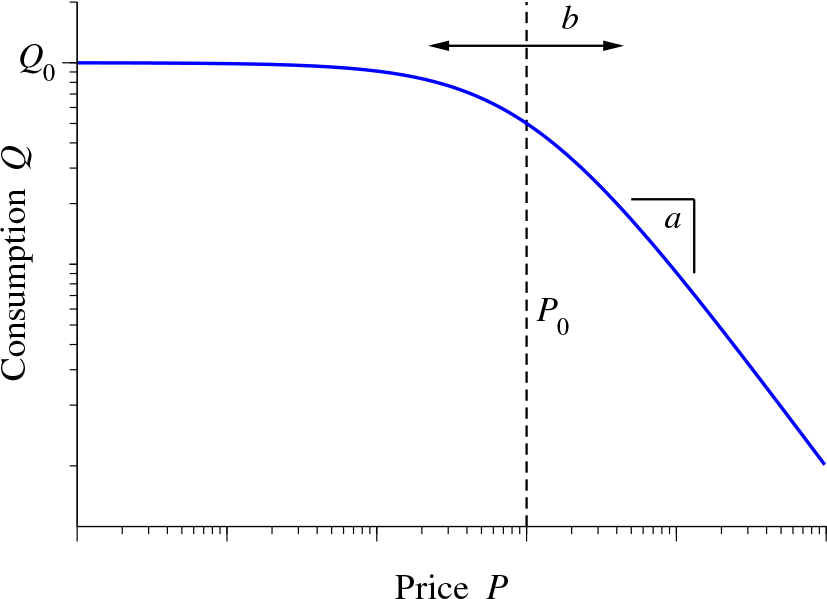
Example of the demand curve form of Eq. (20), along with an indication of the role played by each of the parameters *Q*_0_, *P*_0_, *a*, and *b*.

Figure 4 shows the effect of varying each of these parameters on the shape of the demand curve.

**Fig. 4.**
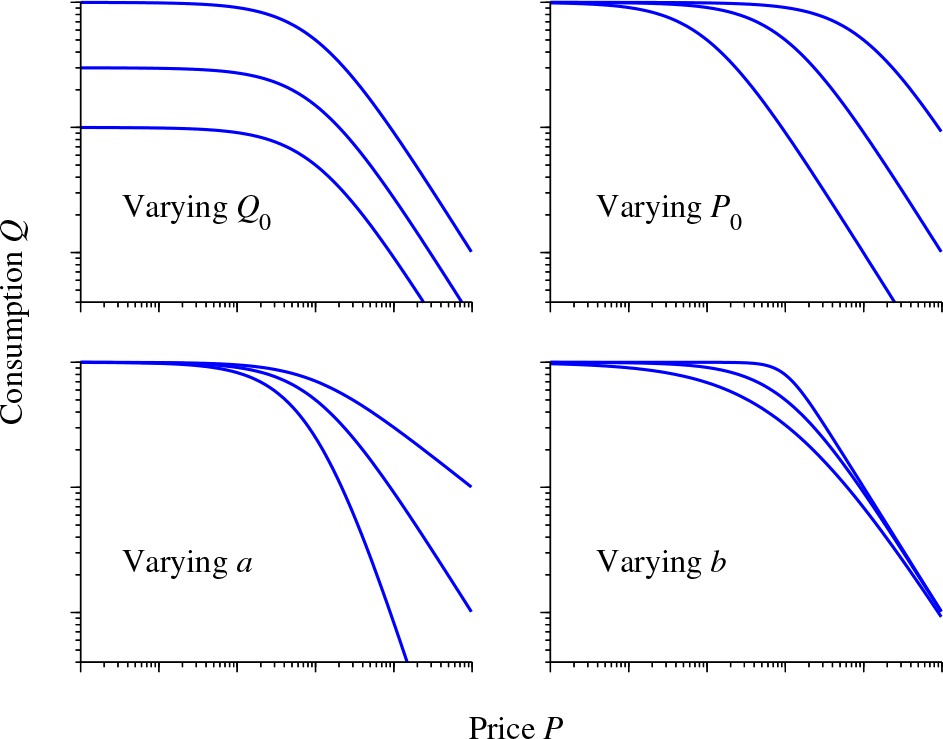
The effect of varying each of the four parameters in Eq. (20).

This form for the demand curve is flexible enough to fit a variety of different data types (see also Figs. 5 and 7). It has three basic regions, one flat, one curved, and one downward sloping, and hence it can fit data with any of these forms, or a combination of all three.

**Fig. 5.**
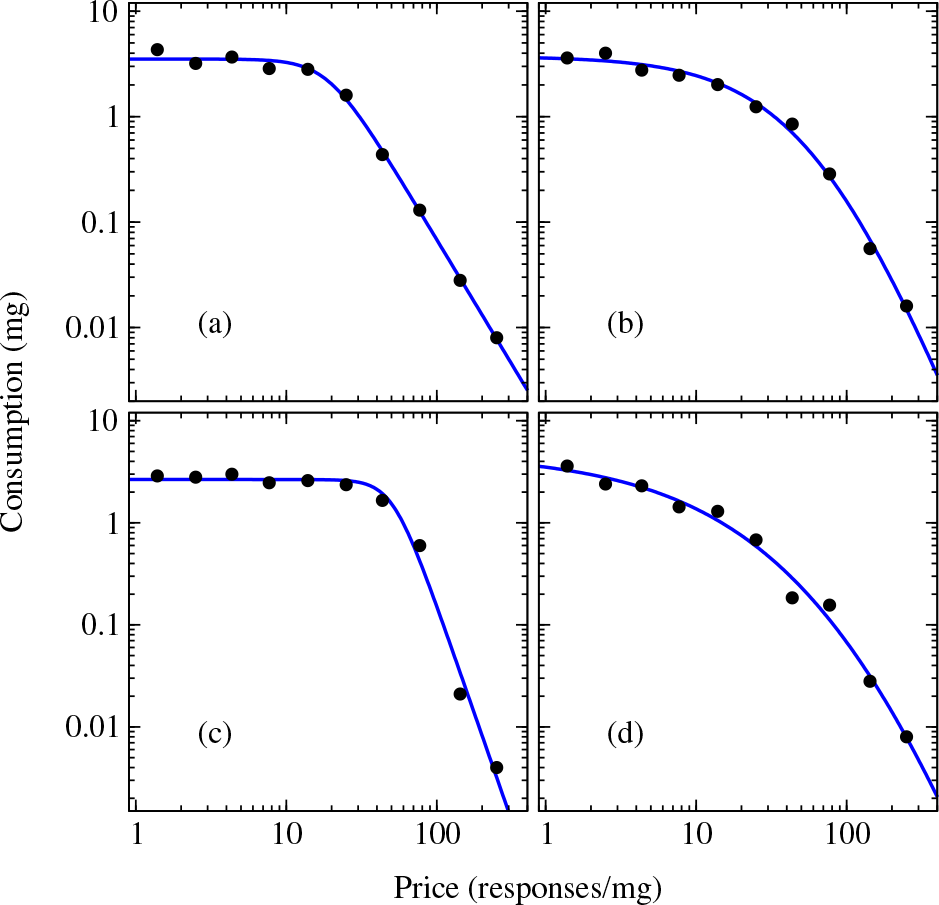
Cocaine consumption as a function of price in four individual male rats. The blue lines show the best fit of the proposed demand curve, Eq. (21).

**Fig. 6.**
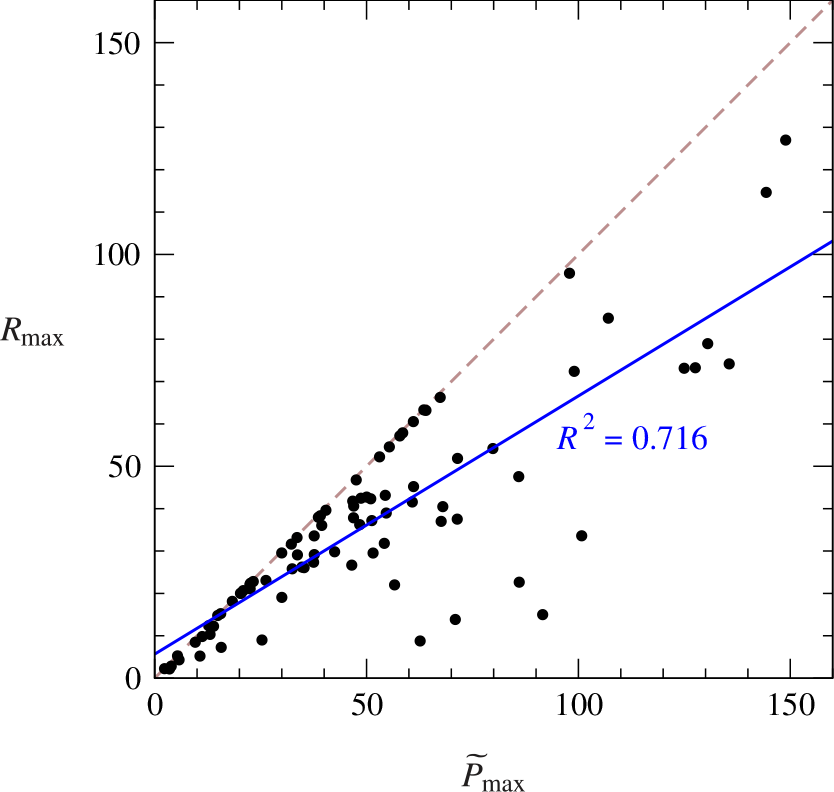
Plot of the values of *R*_max_ against 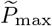 for demand curves for cocaine consumption by nine male rats over 86 separate sessions. The values of the two quantities are substantially correlated (*R*^2^ = 0.716) and clearly obey the rule 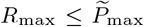 derived at the end of Section 5.2. (The dashed line shows the point at which 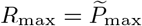, and all points lie on or below this line.) Three extreme outliers on the horizontal axis have been omitted from the plot and from the fit, as discussed in the text.

**Fig. 7.**
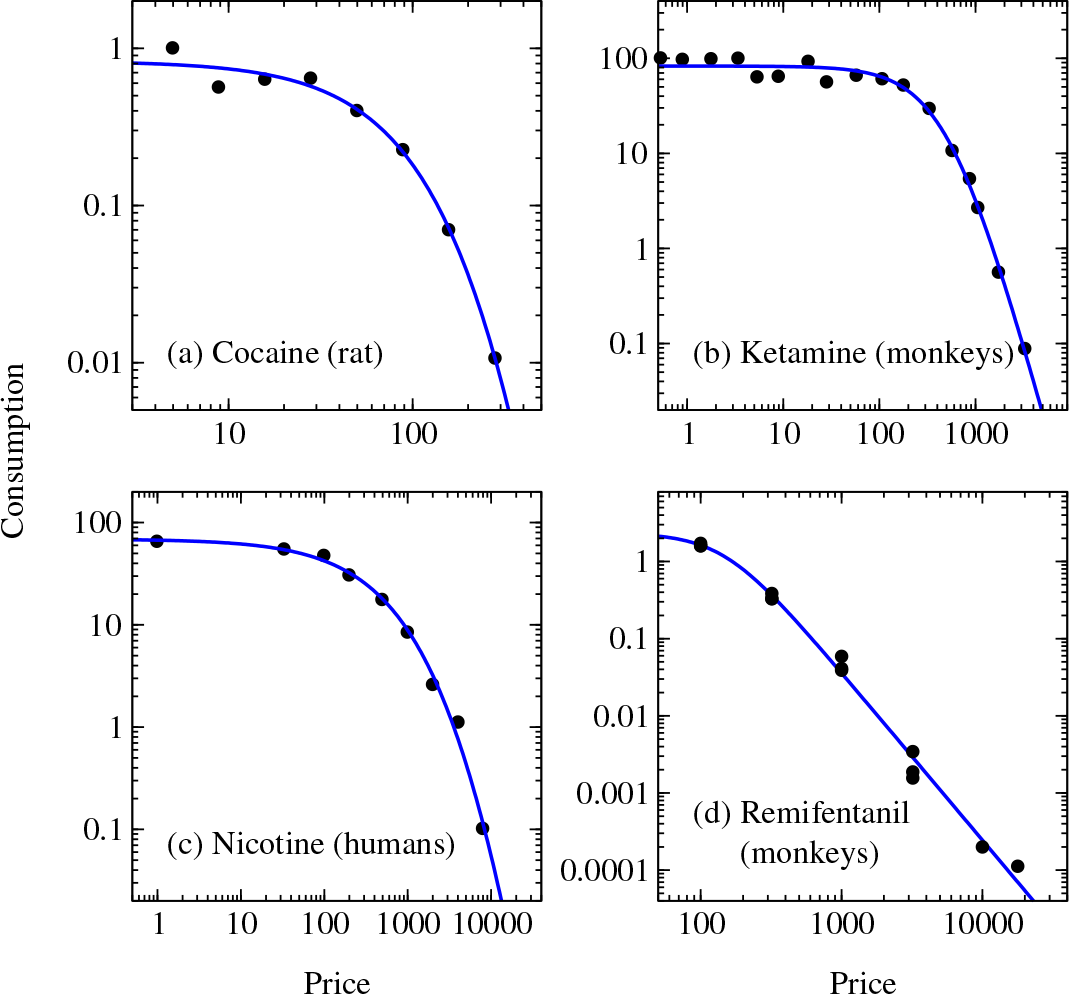
Fits of the demand curve of Eq. (21) to data on self-administration of four different drugs. (a) Cocaine in a male rat from Bentzley et al. (2013), Fig. 3. (b) Ketamine in male monkeys from Winger et al. (2002), Fig. 3, upper panel. (c) Nicotine in humans from Giordano et al. (2001), Fig. 1. (d) Remifentanil in male monkeys from Winger et al. (2006), Table 1.

Since it is common to plot the demand curve on logarithmic scales, one can also rewrite Equation (20) in terms of the log of *Q* thus:

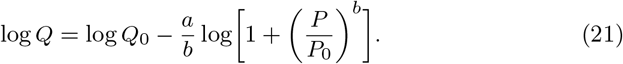

The two forms, Eqs. (20) and (21), are entirely equivalent and contain the same information. One can employ either natural (base *e*) or common (base 10) logarithms— the results are identical either way.

### 5.2 Elasticity and measures of motivation

We can repeat the analyses of Sections 4.3 and 4.4 for this new demand curve. The elasticity *E* can be computed from the logarithmic derivative, Eq. (3), which gives

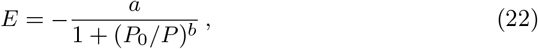

and the maximal value of *E*, the equivalent of Eq. (12), occurs when *P* goes to infinity, giving simply

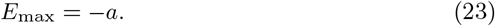

The value of *P*_max_ is given, as previously, by the point at which the elasticity equals −1, i.e., by

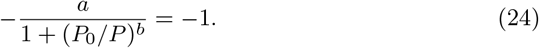

Solving for *P* we find that

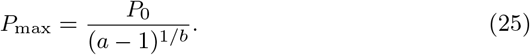

This is a more convenient form than Eq. (14), which involves the special function *W* and requires a complex iterative procedure to calculate *P*_max_ (Hursh and Silberberg 2008). Equation (25) by contrast can be evaluated using only a simple calculator or spreadsheet.

As discussed in Section 3.3, it is useful when comparing values for different goods to normalize the value of *P*_max_ to give a potency- or magnitude-independent measure of motivation. The relevant measure in the present case is

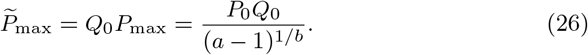

Given the value of *P*_max_ we can compute the corresponding value of *R*_max_ by substituting into Eqs. (4) and (20) to get

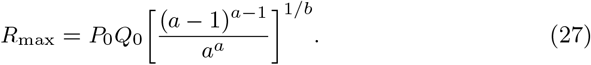

Combining (26) and (27), we can also derive a direct relation between 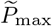 and *R*_max_ thus:

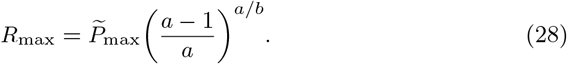

The quantity in brackets is never greater than 1, meaning that we always have 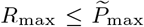, with the equality occurring in the limit where *b* becomes large.^10^ In practice *b* can quite often take values as large as 100 or more, in which case we expect *R*_max_ and 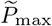 to be essentially equal. These observations shed some light on the question raised in Section 3.2 of the extent to which *R*_max_ and 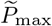 measure the same thing: the answer appears to be that in some cases (but not all) they do, and that overall we expect them to be correlated. We test this conclusion against data in Section 6.

We do not recommend using the values of the fitted parameters *P*_0_, *a*, and *b* themselves as measures of motivation or behavior. When using the exponential form of Eq. (8) [or Eq. (9)] it is common practice, as described in Section 4.4, to use the fitted value of the parameter 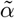 (also denoted *α* elsewhere) as an indicator of reinforcement or motivation, but we do not recommend using the parameters of Eq. (20) in this way since they do not have a clear behavioral interpretation (with the exception of *Q*_0_, which is certainly informative). Instead we recommend using the derived quantities *R*_max_ and 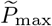.

## 6 Example calculations

Figure 5 shows example fits^11^ of the demand curve of Eq. (21) to data from four different male rats self-administering cocaine under conditions in which each lever press results in an infusion (FR1) and price is varied within session by systematically reducing the dose of drug from 0.72 to 0.004 mg/kg.^12^ As the figure shows, there are a range of different qualitative forms in the data, some with a clear plateau followed by a drop-off, in the classic shape of Fig. 1a, others with a gentler curved form with less of a clear “knee”. Nonetheless the proposed form for the demand curve fits all of the data sets well.

After performing the fits, one can use the fitted parameter values to compute measures such as *P*_max_, 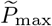, and *R*_max_. The results are shown in Table 3. Note that there is wide variation in the values of both 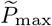 and *R*_max_, and that large values of one do not always correspond to large values of the other. As discussed in Section 5.2, we expect *R*_max_ and 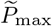 to be correlated, and they must satisfy the constraint 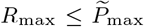, but they are separate measures and can on occasion take widely different values.

**Table 3.** Values of *Q*_0_ for the four fitted curves in Fig. 5 along with the calculated values of the quantities *P*_max_, 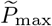, and *R*_max_.

| | $Q_0$ | $P_{\max}$ | $\tilde{P}_{\max}$ | $R_{\max}$ |
| --- | --- | --- | --- | --- |
| a | 3.52 | 17.3 | 60.8 | 41.6 |
| b | 3.74 | 27.0 | 100.8 | 33.6 |
| c | 2.65 | 40.3 | 107.1 | 85.0 |
| d | 4.84 | 19.0 | 91.7 | 15.0 |

To shed more light on the relationship between *R*_max_ and 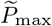, we show in Fig. 6 a plot of the values of *R*_max_ against those of 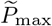 for cocaine self-administration by nine different male rats over ten sessions each, with one point for each session for a total of 86 points (with four sessions omitted for reasons given below). The plot shows substantial correlation between the two measures (*R*^2^ = 0.716) and the fact that 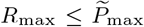 is clear in that all points lie on or below the diagonal dashed line at which 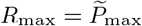. These findings are consistent with previous work examining relationships between *R*_max_ and 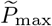 (MacKillop and Murphy 2007; MacKillop et al. 2009).

The results shown in Fig. 6 omit data from four sessions, one for which the demand curve had no point with elasticity −1 (and hence no *P*_max_) and three that returned values of 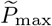 that were extreme outliers, one having a value of almost 500. If these three outliers are included in the fit then most of the correlation disappears (*R*^2^ = 0.155). It is worth asking therefore what causes these outliers. Recall that 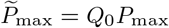 [Eq (25)], so that large 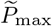 values can be generated either by large *P*_max_ or by large *Q*_0_. We see both behaviors in the present case: one of our three outliers is caused by a large value of *Q*_0_, which appears to be due to the fact that the measured price range failed to span the “knee” in the demand curve, but the other two outliers are caused by large values of *P*_max_. This suggests that 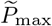 may be a (slightly) less reliable measure of motivation than *R*_max_, which shows no extreme outliers, at least in the data we have examined. This issue would be an appropriate one for further investigation.

Finally, in Fig. 7 we show fits of our proposed demand curve form to data from a range of published studies on consumption of drugs of different drug classes by rats, monkeys, and, in one case, humans. The new form fits this diverse selection of data well, even when the data do not follow the traditional demand curve form as price increases, as in Fig. 7d for example. It is important to note that in such cases the fit will still return values for *P*_max_, *R*_max_, and other quantities but that they may not be meaningful because the price range examined does not cover the plateau and/or knee of the demand curve and hence does not provide a good estimate of *Q*_0_ or *P*_max_. For this reason, it is important to inspect all fits visually, not just for goodness of fit to the data, but also to verify that the behavior of interest is actually captured by the range of prices probed in the experiment.

## 7 Conclusions

The pioneering work of those who brought microeconomics concepts to the study of food and drug self-administration provides a rich foundation on which to build. In this paper we have discussed in detail the use of demand curves and elasticity to quantify the motivating effects of food and drugs in animal experiments. We have highlighted a number of issues with current methodology in this area and proposed a new mathematical framework for the analysis of consumption data that remedies these issues. This framework incorporates a proposed mathematical form for the demand curve that is flexible enough to fit data on a range of reinforcers across different animals and different experimental protocols, provides established metrics of demand (*P*_max_, *R*_max_, *Q*_0_), and the formulas needed to calculate them from fits to data. The equations require only basic algebra for their implementation, although we also provide software implementing our entire analysis pipeline, including data fits, plotting, and the calculation of summary statistics (see appendix).

## Acknowledgements

The authors thank Crystal Carr, Alex B. Kawa, Lauren Longyear, and Terry E. Robinson for providing the data used in Figs. 1, 2, 5, and 6 and Terry E. Robinson for useful comments during the preparation of this manuscript. This work was supported in part by the National Institutes of Health under grants NIDA R01DA044204, R21DA045277, T32DA007268, and NIDDK R01DK106188.

## Author Contributions

The research was designed jointly by MEJN and CRF. MEJN performed mathematical analysis and wrote the manuscript. CRF performed data analysis and wrote the manuscript.

## A Analysis software

In the accompanying online materials (http://umich.edu/~mejn/demandcurve) we provide a software program that performs the analyses described in this paper using the proposed demand curve of Eq. (20). In this appendix we describe the use of this program. The same information can also be found in the form of a video tutorial, also in the accompanying materials.

### A.1 Installation

The program is called demand.py. To install it, simply download it from the web address given above and place it in the folder or directory containing the data you want to analyze.

The program is written in the Python computer language. Running it requires that you also have the Python language installed. Many computers come with Python already installed, but if yours does not, Python is free to download from the web. There are various versions available, but we recommend the “Anaconda” Python distribution, which contains everything you will need in one single package. Anaconda is available for free from www.anaconda.com. Two versions are currently available, version 2 (actually version 2.7 at the time of writing) and 3 (version 3.7 at the time of writing). Our program will work with either but we recommend installing version 3 simply because it is the most up-to-date.

If you wish to install Python by another route (for instance using a package manager), then you should, at a minimum, install the Python language itself (either version 2 or version 3), plus the packages “scipy” (for curve fitting) and “matplotlib” (for graphics).

### A.2 Data input

The program takes input data in the form of prices and responses. It can handle an arbitrary number of price points in a single analysis and can analyze data from multiple animals or multiple sessions in a single run. Data should be prepared in a spreadsheet as shown in Fig. 8. The first column of the sheet contains the prices, in any units you choose. Prices can be in any order—they need not be in increasing (or decreasing) order, though they can be. The second and subsequent columns contain the response at each price in terms of the amount of work (e.g., number of lever presses) done in experimental sessions, with one column for each session or data set. There can be just a single session (for a total of two columns—prices and responses) or many (for a total of *n* + 1 columns if there are *n* sessions). There should be no blank cells, nor empty rows or columns, in the spreadsheet, except for the unused rows and columns below and to the right of the data. (If you wish to analyze data from experiments using different numbers of price points then you will have to use separate spreadsheets.)

**Fig. 8.**
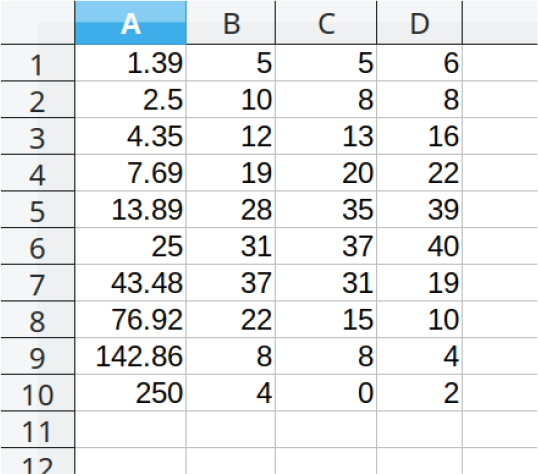
Example of the format of the input data. The first column (column A) represents prices, which can be in any units. (Ours are in responses per milligram.) Second and subsequent columns, of which there can be any number, represent responses at each price point for separate animals, sessions, or data sets. In this case the file has columns for three different sessions (columns B, C, and D). Our responses are whole numbers, since they represent lever presses by individual animals, but one can also use responses averaged over several animals or sessions, in which case the values need not be whole numbers. When complete, the spreadsheet should be saved in CSV (comma-separated value) format with commas as separators between the fields.

The spreadsheet should be saved in CSV (comma-separated value) format, using commas as separators between the values and the file extension “.csv”. (Note that CSV files can be saved with spaces as separators, but this should be avoided as it will not work with our program.) When using Microsoft Excel, for instance, saving as a CSV file is a standard option under the “Save as” menu item; any of the sub-formats listed there (Mactintosh, MS-DOS, or CSV) will work. An example input file, called `example.csv`, is included in the accompanying online materials.

### A.3 Running the program

#### Windows

On an appropriately configured Windows system it will be possible to run the program from a command prompt window (also known as a “DOS window”), by changing to the folder where the program file resides and typing either “py demand.py” or “python demand.py” into the command window (depending on how the computer is set up). Alternatively, the program can be run inside a Python development environment such as Jupyter, Idle, or Spyder.

The latter all are included, for instance, in the Anaconda distribution mentioned above. When Anaconda is started it will display a screen offering a choice of environments for use. Start the environment of choice by double-clicking on its launch icon, then load the program demand.py and run it. Specific procedure will depend on which environment you use. See the video tutorial for an example using the Spyder environment.

#### OSX

On a Mac one can normally open a command window and type “python demand.py” to run the program, or use any of the several available Python development environments—see the instructions for Windows users above.

#### Linux

Under Linux one can run the program from the command line with the command “python demand.py” or from within a development environment.

### A.4 Using the program

When first run, the program will ask for the name of the input data file. Type in the name of the CSV file containing your data. (The name is case-sensitive, so take care with capitalization.) The file extension “`.csv`” at the end of the name is optional and can be omitted. The file name can also be provided to the program as a command-line argument when the program is run. For instance, under Windows one might type “`py demand.py example.csv`” and the program would use the data file `example.csv`. This is a convenient feature when running the program in batch mode (i.e., unattended).

Once the file name is entered, the program will perform the analysis. It works by computing consumption levels from the response data then performing a nonlinear least-squares fit of the logarithms of the consumption values to Eq. (21). The output of a typical run of the program looks like this:

~~~
Data read from file example.csv
  Price points: 10
  Curves: 3
Analyzing curve 1
Analyzing curve 2
Analyzing curve 3
~~~

The program gives a summary of the data it found in the input file (number of price points and number of data sets), then lists the demand curves one by one as it analyzes them. The program may also print out cautionary messages if it believes there are potential problems with the data. For instance, if the fitted value of *Q*_0_ lies well outside the range of observed consumption it will print a message like this:

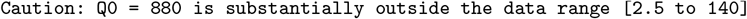

This could be an indication that the experiment failed to probe the salient portion of the demand curve near the breakpoint at which demand falls off.

When the program finishes, which typically takes a few seconds, it will produce a set of output files in the same folder. First, there will be an output data file. If the input file was called `example.csv`, the output file will be called `example_params.csv`. This is a CSV spreadsheet containing the best fit values of the parameters *Q*_0_, *P*_0_, *a*, and *b* for each session in the input file, plus the values of the derived quantities *R*_max_, *P*_max_, and 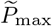. This file can be opened in Microsoft Excel, Google Docs, or any similar spreadsheet program. There is one row of the file for each session, in the same order as the columns of the input file, so that the row for curve 1 corresponds to data in column B of the input file, curve 2 to data in column C, and so forth. An example output file is shown in Fig. 9.

**Fig. 9.**
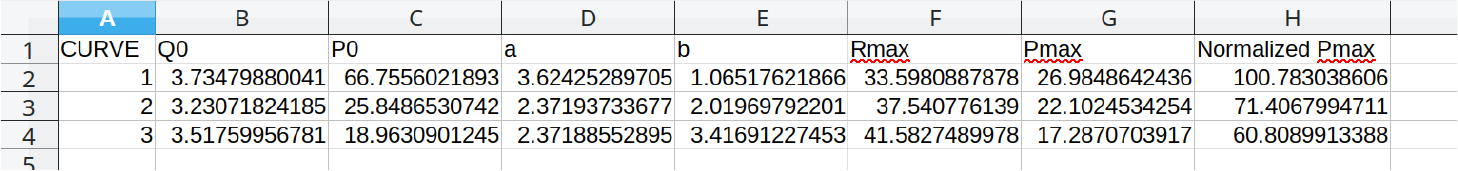
Example of the output spreadsheet generated by the program. Columns represent the parameters *Q*_0_, *P*_0_, *a*, and *b* of the fitted demand curves, plus the derived values *R*_max_, *P*_max_, and 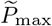 (“Normalized *P*_max_”), as indicated, and there is one row of the spreadsheet for each data set in the input file.

Second, the program produces a set of output figure files, one for each input session or data set (i.e., for each column from B onward in the original input). An example is shown in Fig. 10. Each figure file contains two graphs: the left graph shows, as a function of price, the data points for consumption in blue and the fitted demand curve in green; the right graph shows the original data for responses in blue and the fitted curve in green. If the input data file is called `example.csv` then these figure files will have names `example 1.png`, `example 2.png`, and so forth, numbered in the same order as the columns of the input file.

**Fig. 10.**
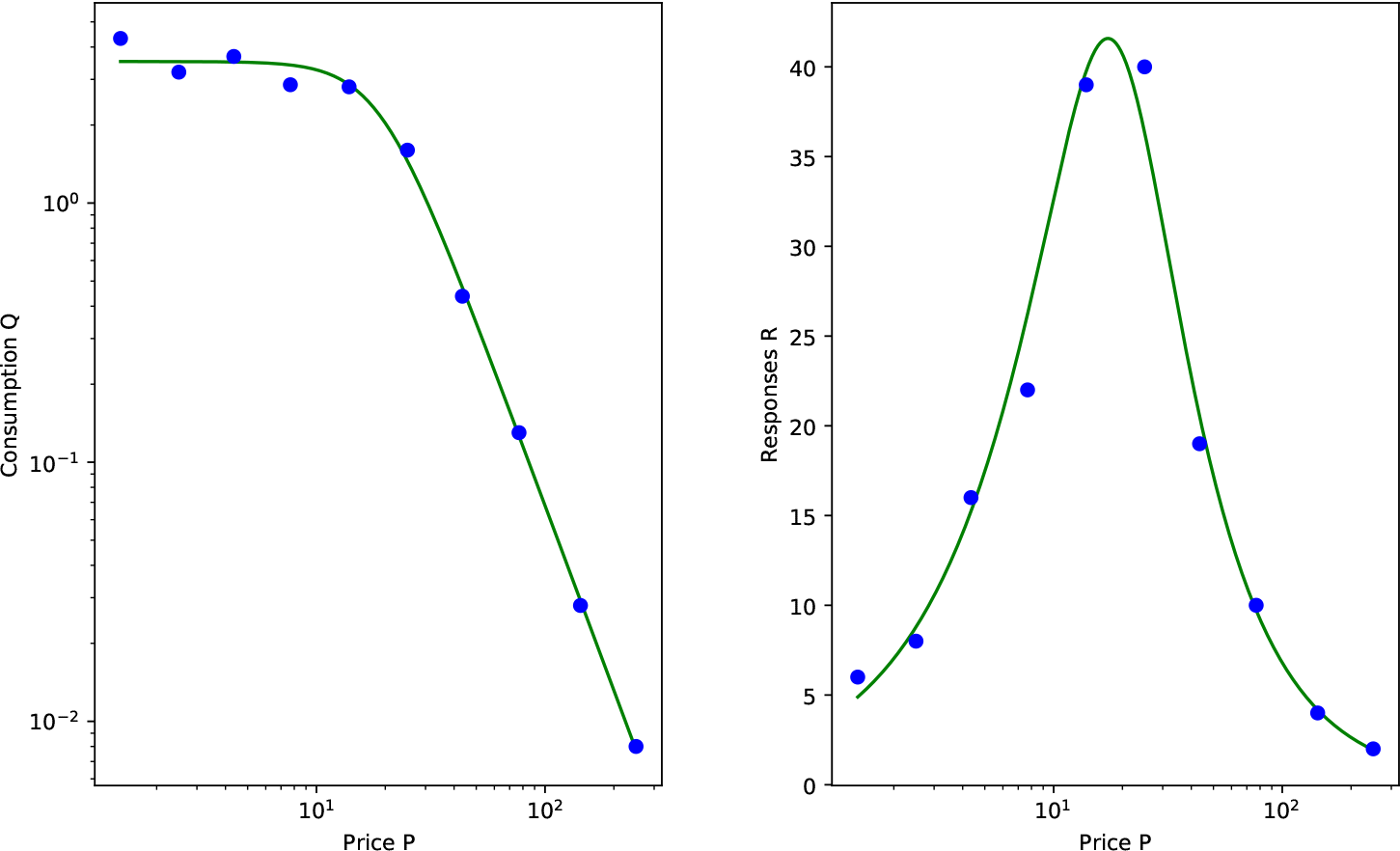
A typical figure produced by the program. Left: data for consumption against price (points) and the best fit of the demand curve from Eq. (20) as determined by the program (solid line). Right: the original response data against price (points) and the corresponding revenue curve determined from the demand curve using Eq. (4) (solid line). *R*_max_ is the highest value of the revenue curve and *P*_max_ is the price at which that highest value occurs. In this case we find *R*_max_ = 41.6 responses and *P*_max_ = 17.3 responses per milligram.

It is a good idea before using the numerical results of any of the fits to inspect the corresponding figures to make sure that the fit is a reasonable one. If the data fluctuate a lot, or if the experiment fails to probe the correct price range, then the fit may be a poor one. In cases where the data are entirely unlike the expected demand-curve form of plateau-plus-decline, the fit may fail and the fitted curve will not resemble the data points. Some thought may be needed to interpret the results of the procedure in any specific case.

## Footnotes

1 Previously unpublished data kindly provided by A. B. Kawa, L. Longyear, and T. E. Robinson.

2 Technically, the derivative is the limiting value of *dQ/dP* for infinitesimal *dP* and *dQ*.

3 The notation *R* for revenue is a standard one in economics, but in the animal literature one sometimes sees revenue denoted *O* for “output.” See also Table 1.

4 Sometimes also denoted *O*_max_.

5 Some authors have employed common logarithms (base 10) instead, which case some of the formulas and parameter values are modified, but the scientific outcome is unchanged.

6 The value of *k* must also always be greater than the mathematical constant *e* = 2.718 for *P*_max_ to be well defined (Gilroy et al. 2019). If *k* is smaller than this then there is no point on the demand curve where elasticity is −1 and hence there is no *P*_max_—see Section 3.2.

7 Some authors have used common (base 10) logarithms instead of natural (base *e*) logarithms, in which case the solution takes a slightly different form—see Gilroy et al. (2019).

8 Price in this context is often written *C* rather than *P*. This, however, is merely a notational choice.

9 See http://umich.edu/~mejn/demandcurve for details.

10 The same result also holds for the exponential demand curve. Equation (15) can be rewritten as 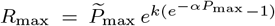, but 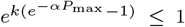 always, so this implies that 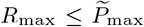.

11 We perform our fits using a least-squares best fit of Eq. (21) to the logarithm of consumption, discarding data points with zero consumption, since one cannot take the log of zero. It is also possible to fit consumption data directly to Eq. (20) and it is a straightforward modification of the approach to do so, but in practice we find this gives visually poorer fits. See also Koffarnus et al. (2011).

12 Unpublished data kindly provided by C. Carr and T. E. Robinson.

## References

Aston ER, Cassidy RN (2019) Behavioral economics assessments in the addictions. Current Opinion in Psychology 30:30–42

Ayers RM, Collinge RA (2004) Microeconomics. Prentice Hall, Upper Saddle River, NJ

Becker GS, Grossman M, Murphy KM (1994) An empirical analysis of cigarette addiction. American Economic Review 84:84–396

Bentzley BS, Fender KM, Aston-Jones G (2013) The behavioral economics of drug self-administration: A review and new analytical approach for within-session procedures. Psy-chopharmacology 226:226–113

Bickel W, Marsch L, Carroll M (2000) Deconstructing relative reinforcing efficacy and situating the measures of pharmacological reinforcement with behavioral economics: A theoretical proposal. Psychopharmacology 153:153–44

Cartwright E (2018) Behavioral Economics, 3rd edn. Routledge, New York

Galuska C, Banna K, Willse L, Yahyavi-Firouz-Abadi N, See R (2011) A comparison of economic demand and conditioned-cued reinstatement of methamphetamine-seeking or food-seeking in rats. Psychopharmacology (Berl) 22(4):312–323

Gilroy SP, Kaplan BA, Reed DD, Hantula DA, Hursh SR (2019) An exact solution for unit elasticity in the exponential model of operant demand. Experimental and Clinical Psychopharmacology March 28:advance publication online, URL http://dx.doi.org/10.1037/pha0000268

Giordano LA, Bickel WK, Shahan TA, Badger GJ (2001) Behavioral economics of human drug self-administration: Progressive ratio versus random sequences of response requirements. Behavioural Pharmacology 12:12–343

Hursh SR (1993) Behavioral economics of drug self-adminstration: An introduction. Drug and Alcohol Dependence 33:33–165

Hursh SR, Silberberg A (2008) Economic demand and essential value. Psychological Review 115:115–186

Hursh SR, Winger G (1995) Normalized demand for drugs and other reinforcers. Journal of the Experimental Analysis of Behavior 64:64–373

Hursh SR, Galuska CM, Winger G, Woods JH (2005) The economics of drug abuse. Molecular Interventions 5:5–20

Kahneman D (2011) Thinking Fast and Slow. Farrar, Straus and Giroux, New York

Kawa A, Bentzley B, Robinson T (2016) Less is more: prolonged intermittent access cocaine self-administration produces incentive-sensitization and addiction-like behavior. Psychopharmacology (Berl) 233:233–19

Koffarnus MN, Hall A, Winger G (2011) Individual differences in rhesus monkeys’ demand for drugs of abuse. Addiction Biology 17:17–887

Lacy RT, Austin BP, Strickland JC (2019) The influence of sex and estrous cyclicity on cocaine and remifentanil demand in rats. Addiction Biology pp 1–10

MacKillop J, Murphy JG (2007) A behavioral economic measure of demand for alcohol predicts brief intervention outcomes. Drug and Alcohol Dependence 89:89–227

MacKillop J, Murphy JG, Tidey JW, Kahler CW, Ray LA, Bickel WK (2009) Latent structure of facets of alcohol reinforcement from a behavioral economic demand curve. Psychopharmacology 203:203–33

Oleson EB, Richardson JM, Roberts DC (2011) A novel iv cocaine self-administration procedure in rats: differential effects of dopamine, serotonin, and gaba drug pre-treatments on cocaine consumption and maximal price paid. Psychopharmacology 214:214–567

Olver FWJ, Lozier DM, Boisvert RF, Clark CW (2010) NIST Handbook of Mathematical Functions. Cambridge University Press, Cambridge

Pantazis C, James M, Bentzley B, Aston-Jones G (2019) The number of latera-lhypothalamus orexin/hypocretin neurons contributes to individual differences in cocaine demand. Addiction Biology Jul 11:[Epub ahead of print]

Perloff JM (2016) Microeconomics: Theory and Applications with Calculus, 4th edn. Pearson, New York

Richardson N, Roberts D (1996) Progressive ratio schedules in drug self-administration studies in rats: a method to evaluate reinforcing efficacy. Journal of Neuroscience 66:66–1

Winger G, Hursh SR, Casey KL, Woods JH (2002) Relative reinforcing strength of three N-methyl-D-aspartate antagonists with different onsets of action. Journal of Pharmacology and Experimental Therapeutics 301:301–690

Winger G, Galuska C, Hursh S, Woods J (2006) Relative reinforcing effects of cocaine, remifentanil, and their combination in rhesus monkeys. Journal of Pharmacology and Experimental Therapeutics 1318:223–229

